# A Transgenic System for Targeted Ablation of Reproductive and Maternal-Effect genes

**DOI:** 10.1101/2020.10.22.351403

**Authors:** Sylvain Bertho, Odelya Kaufman, KathyAnn Lee, Adrian Santos-Ledo, Daniel Dellal, Florence L. Marlow

## Abstract

Maternally provided gene products regulate the earliest events of embryonic life, including formation of the oocyte that will develop into an egg, and eventually an embryo. Forward genetic screens have provided invaluable insights into the molecular regulation of embryonic development, including essential contributions of some genes whose products must be provided to the transcriptionally silent early embryo for normal embryogenesis, maternal-effect genes. However, other maternal-effect genes are not accessible due to their essential zygotic functions during embryonic development. Identifying these regulators is essential to fill the large gaps in our understanding of the mechanisms and molecular pathways contributing to fertility and maternally regulated developmental processes. To identify these maternal factors, it is necessary to bypass the earlier requirement for these genes so that their potential later functions can be investigated. Here we report reverse genetic systems to identify genes with essential roles in reproductive and maternal-effect processes, as proof of principal and to assess the efficiency and robustness of mutagenesis we used these transgenic systems to disrupt two genes with known maternal-effect functions, *kif5Ba* and *bucky ball*.

**Summary Statement:** We report reverse genetic systems to identify essential regulators of reproductive and maternal-effect processes, as proof of principal we used these transgenic systems to disrupt genes with known maternal-effect functions.

## Introduction

Maternally provided gene products regulate the earliest events of embryonic life, including formation of the oocyte that will develop into an egg, and eventually an embryo. Disruption of oocyte development or early embryonic cleavages and patterning cause congenital birth defects and are apparent in 2-5% of human births according to sources such as the National Institute of Child Health and Human Development. For some of these birth defects chromosomal aneuploidy is causally associated, but for others the genetic and molecular basis remains poorly defined (Ambartsumyan and Clark, 2008; Hassold et al., 2007). The most devastating mutations to the embryo are those occurring in early embryogenesis at stages when the diverse cell types necessary to build an embryo are being specified and the basic body plan is being laid down. Mutations occurring in later development are detectable in the children born with the consequent birth defects. In contrast, mutations in genes essential for processes occurring before implantation or during gastrulation are more severe and result in embryonic lethality (Ambartsumyan and Clark, 2008; Hassold et al., 2007). In mammals, embryonic development occurs *in utero;* so, mutations disrupting essential regulators of early embryogenesis often go undetected due to arrest *in utero* and miscarriage (Hassold et al., 2007; Hassold and Hunt, 2007; Zhao et al., 2006). Consequently, our understanding of the molecular and genetic regulation of this extremely sensitive developmental period remains incomplete.

In fish and humans, the earliest developmental events, are regulated by maternally supplied gene products because the early zygote is transcriptionally silent (Abrams et al., 2020; Marlow, 2010; Sato et al., 2019; Vastenhouw et al., 2019; Wu and Vastenhouw, 2020; Zhao et al., 2006). These maternally supplied genes are known as maternal-effect (ME) genes because the embryo relies on gene function supplied by its mother. Mutations disrupting these genes in the mother will result in phenotypically abnormal progeny regardless of the embryo’s genotype. Where studied, the basic aspects of oocyte development are remarkably conserved among animals, including regulation of meiotic initiation and arrest (pauses at specific stages in the meiotic cycle, reviewed in (Marlow, 2010). In recent years a number of genes with maternal functions have been discovered in the mouse; however, the precise contribution of ME genes is often masked by the *in utero* arrest of the mutant embryos (Marlow, 2010; Wu and Dean, 2020; Zhao et al., 2006). That is, while it is clear the gene is essential, in some cases the phenotype is detected solely based on a lack of or underrepresentation of mutant genotypes among the progeny (if any) of mutant mothers; so, determining why and when the embryo arrests is hampered by the low recovery of mutant progeny. The external fertilization and development of the zebrafish and the large numbers of progeny produced each week make it possible to recover and examine every egg that is produced and to pinpoint the cellular and molecular basis of the developmental defect or embryonic lethality.

Forward genetic screens have provided invaluable insights into the molecular regulation of embryonic development, including essential contributions of some maternal-effect genes (Dosch et al., 2004; Pelegri et al., 2004; Pelegri and Mullins, 2004; Wagner et al., 2004). However, any maternal-effect gene with an additional essential zygotic function during embryonic development cannot be identified in this way because the mutants do not reach reproductive maturity. Indeed, although the zebrafish is an excellent genetic system, traditional mutagenesis strategies and modern reverse genetic approaches alone have not allowed for straightforward identification of maternal functions of zygotic lethals (Doyon et al., 2008; Foley et al., 2009a; Foley et al., 2009b; Lawson and Wolfe, 2011; Moens et al., 2008; Sander et al., 2011a; Sander et al., 2011b; Sood et al., 2006). To identify these factors, it is necessary to bypass the early zygotic requirement for the gene so that potential reproductive or maternal functions of the gene can be investigated. Methods to circumvent these zygotic lethal phenotypes in the zebrafish were pioneered in the Schier and Raz labs (Ciruna et al., 2002). Their germline replacement approach takes advantage of the early separation of somatic and germline lineages in zebrafish to generate animals with a normal somatic composition and a mutant germline through host germline ablation and transplantation (replacement) with mutant donor germ cells (Ciruna et al., 2002). This strategy allows the animal to survive to produce mutant gametes, which can be examined for reproductive and maternal-effect phenotypes. Although this approach has been applied to examine the function of specific genes (Bennett et al., 2007; Borovina et al.; Ciruna et al., 2006; Ciruna et al., 2002; Williams et al., 2018), thus far, no systematic germline replacement screen of the existing large collection of zebrafish zygotic lethal mutations has been attempted because this approach is tedious and germline replacement is inefficient.

One drawback of the current strategy to investigate genes with ME functions, *dead end* mediated germline replacement, is that few females are produced using this approach (Ciruna et al., 2006; Ciruna et al., 2002). This is a clear impediment to studies of oocyte development or ME gene functions. It is currently thought that this male bias is in part due to insufficient numbers of donor PGCs to support female specific gonadogenesis as *dead end* morpholino depleted embryos develop as sterile males (Ciruna et al., 2002; Siegfried and Nusslein-Volhard, 2008; Slanchev et al., 2005; Weidinger et al., 2003). Additional evidence for a signal from the germline (e.g. oocytes) to support female gonad development and maintenance comes from the phenotypes of zebrafish mutants in which oocyte survival is compromised (Cao et al., 2019; Dranow et al., 2016; Dranow et al., 2013; Hartung et al., 2014; Rodriguez-Mari et al., 2010; Rodriguez-Mari and Postlethwait, 2011; Rodriguez-Mari et al., 2011). In these mutants a lack of oocytes results in failure to antagonize masculinizing signals and leads to female to male sex reversal (Rodriguez-Mari et al., 2010; Rodriguez-Mari and Postlethwait, 2011; Rodriguez-Mari et al., 2011). To circumvent this problem, we utilized a transgenic mutagenesis approach to generate mosaic gonads in which germ cells carrying the mutagenic cassettes and potential mutations are marked with fluorescent reporters.

Here we report reverse genetic systems to identify genes with reproductive and maternal-effect functions. This approach will be particularly useful for genes whose maternal-effect functions are masked by their essential zygotic roles in early embryogenesis. This is possible because this transgenic approach selectively mutates the germline and thus allows the animal to survive to produce mutant gametes, which can be examined for spermatogenesis, oogenesis, egg, and early embryo phenotypes. As proof of concept we used this system to attempt to phenocopy two genes known to have adult reproductive and maternal-effect phenotypes, *kif5Ba* and *bucky ball*. The potential to examine the function of every gene in its genome makes the zebrafish an extremely powerful vertebrate system to unravel molecular and genetic control of developmental processes and adult physiology and disease. Complete phenotypic characterization of the zebrafish mutants, or determination of the zebrafish phenome, will significantly improve our understanding of processes that are difficult to access in mammals, in particular maternal-effect processes.

## MATERIALS AND METHODS

### Animals

Mutant fish strains were generated using Crispr-Cas9 mutagenesis with modifications (see the plasmids list, Supplemental Table 1) to the plasmid backbone published in (Ablain et al., 2015). All procedures and experimental protocols were performed in accordance with NIH guidelines and were approved by the Einstein (protocol #20140502) and Icahn School of Medicine at Mount Sinai Institutional (ISMMS) Animal Care and Use Committees (IACUC #2017-0114).

### Primers

All primers are listed in Supplemental Table 2.

### OMS and GMS mutagenesis plasmids

*oms* and *gms* plasmids (Supplemental Table 1) were created using the tissue-specific promoter system described in Ablain et al (Figure 1, Supplemental Methods). In brief, we digested a p3E_polyA_U6:gRNA (Figure 1B, Supplemental Methods) (Ablain et al., 2015) using BseRI enzyme and then inserted annealed gene-specific gRNA targeting *kif5Ba* and *buc*. For the gRNAs, previously validated gRNA sequences were used to target *kif5Ba* (Campbell et al., 2015), *rbpms2* (Kaufman et al., 2018) and to target *buc*, new gRNAs were tested for mutagenic activity (Supplemental Table 2). Gateway recombination reactions were then used to generate expression constructs with Cas9 driven by the *bucky ball* (Heim et al., 2014) or *ziwi* (Leu and Draper, 2010) promoter. Recombination order was confirmed by sequencing using *ziwi* promoter or *buc* promoter, *cas9* and *U6* promoter primers (Supplemental Table 2). The complete procedure is detailed in the Supplemental Methods.

**Figure 1.**
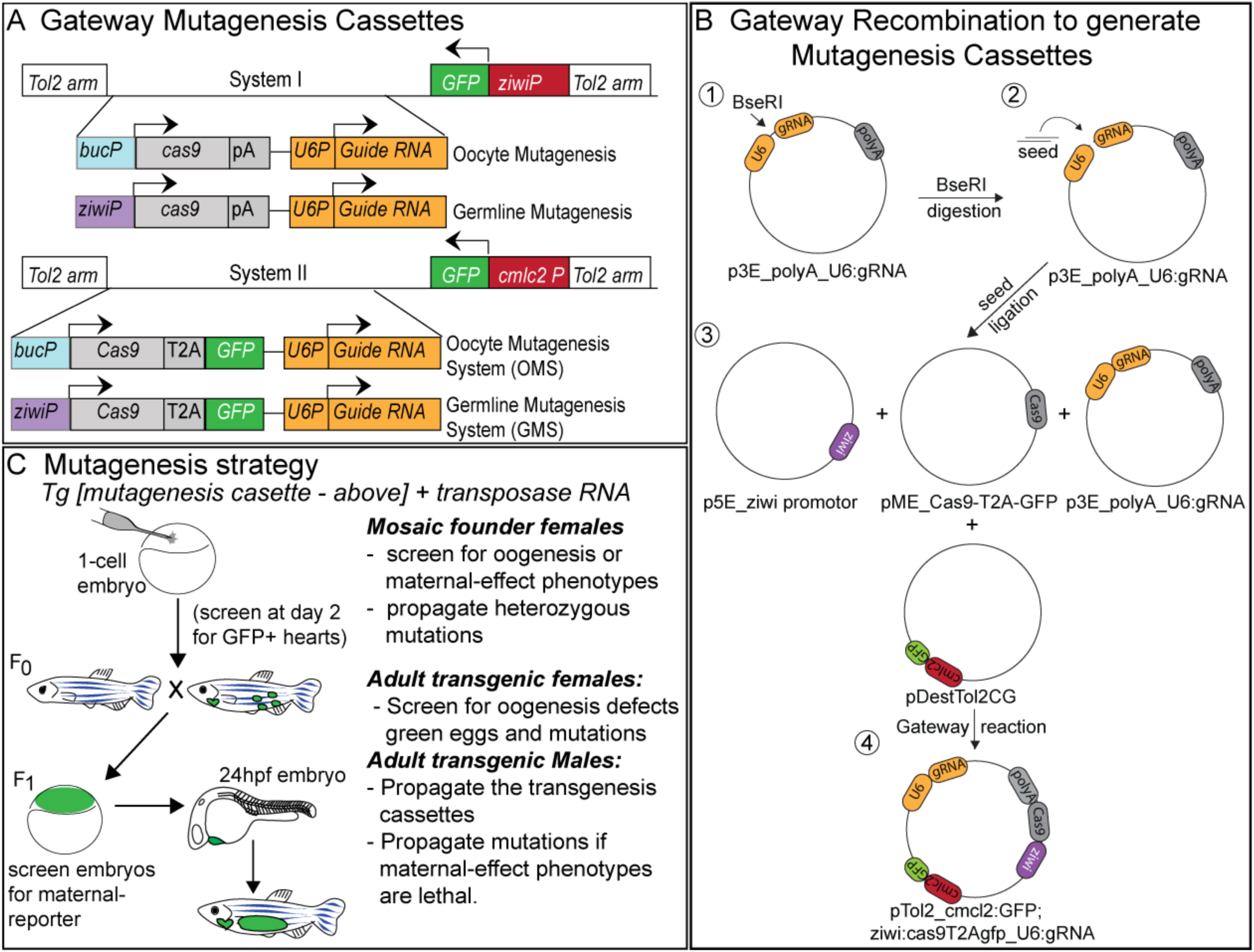
Ovary and Germ Cell Mutagenesis Systems. A) Schematic depicting the gateway mutagenesis system for germline induced CRISPR/Cas9 mutagenesis. System I vectors express Cas9 protein from either the *bucky ball* (oocytes) or *ziwi* (all germ cells) promoter. Target specific guide RNAs are expressed from the U6 promoter, and GFP is expressed from the *ziwi* promoter to mark all germ cells with the vector. System II vectors are similar except Cas9 is expressed as a single transcript with GFP following a T2A cleavage signal. This self-cleaving peptide marks cells expressing Cas9 protein. In addition, GFP is expressed from an independent somatic cell promoter, *cardiac myosin light chain 2 (cmlc2)*, to allow for selection of transgenic positive animals at 2dpf, based on expression of GFP in the heart. B) Schematic of vectors and gateway mediated recombination to generate *oms* and *gms* mutagenesis plasmids. C) Schematic of the mutagenesis strategy. Briefly, mutagenesis cassette and transposase are injected to the one-cell stage embryo. For system II putative founders are selected for expression of GFP in the heart at 2dpf. The gonads can be screened directly for germline mutations, or the F1 progeny of founders can be screened for maternal *(buc* or *ziwi* promoter) or paternal-effect *(ziwi* promoter) mutations/phenotypes.

### Stable transgenic lines

To generate stable *oms* and *gms* transgenic lines, Tol2 Transposase RNA was transcribed from pCS2FA-transposase (Kwan et al., 2007), and combined with *oms* or *gms* vector circular DNA (25 ng/μl each). Embryos were injected with 1 nl of the plasmid/transposase solution at the one-cell stage. Embryos with GFP positive hearts were selected at 2 dpf and raised to generate founders.

### Genotyping

Genomic DNA was extracted from adult fins using standard procedures (Meeker et al., 2007). The genomic region surrounding *kif5Ba* was amplified using the primers 5’-GGAGTGCACCATTAAAGTCATGTG −3’ and 5’-GTCGGTGTCAAATATTGAGGTC-3’. The genomic region surrounding *buc* was amplified using the primers 5’-TGCAGTATCCTGGCTATGTGAT-3’ and 5’-ACCACATCAGGGGTAGAAGAGA-3’. The genomic region surrounding *rbpms2* was amplified using primers 5’-GGGAAGCACCGCTTACAATA-3’ and 5’-TTTGACTCACATGGGTCTCG-3’ (Supplemental Table 2) After 35 cycles of PCR at 59°C annealing for *kif5Ba* and 57°C for *buc*, PCR fragments were TA cloned into pCR4-TOPO vector (K457502, Invitrogen). After transformation, each plasmid was checked by sequencing using universal primers.

### Western blot

25 transgenic (identified by GFP expression) or not transgenic (GFP negative) were pulled together at 2 dpf. Embryos were pipetted up and down in PBS and centrifuged for one minute at 3000 rpm. Supernatant was discarded and pellet was resuspended in 50μl SDS 2x Buffer. 10μl per sample was loaded in a 12% SDS-page gel and proteins were transferred to PVZ membranes. Anti-CRISPR antibody (Abcam, ab204448). Membranes were imaged in G:BOX Chemi XX6 (Syngene).

### *In situ* hybridization

For *in situ* hybridization, embryos at the specified stages, were fixed in 4% paraformaldehyde overnight at 4°C. *in situ* hybridization was performed according to (Thisse et al., 2004), except hybridization was performed at 65°C. In addition, maleic acid buffer (100mM maleic acid, pH 8, 150mM NaCl) was substituted for PBS during the antibody incubations, and BM Purple was used to visualize the RNA probes (Roche, 1442074).

### Immunostaining and Imaging

For whole-mount immunofluorescence stained shield, 30hpf embryos, or ovaries, tissues were fixed in 3.7% paraformaldehyde overnight at 4°C. The following day the samples were washed in PBS, dehydrated by washing in MeOH, and then stored at −20°C. To visualize germ cells, chicken anti-GFP antibody (Invitrogen, A10262) was used at a 1:500 dilution. Secondary antibodies: Alexafluor488, CY3, (Molecular Probes) were diluted 1:500. Samples were mounted in vectashield with DAPI and images were acquired using a Zeiss Axio Observer inverted microscope equipped with Apotome II and a CCD camera, a Zeiss Zoom dissecting scope equipped with Apotome II. Image processing was performed in Zenpro (Zeiss), ImageJ/FIJI, Adobe Photoshop and Adobe Illustrator.

## RESULTS AND DISCUSSION

### Vector based system for generation of germline and maternal-effect mutants

To date traditional mutagenesis strategies and modern reverse genetic approaches alone have only provided limited access to maternal functions of genes with essential zygotic functions in zebrafish (Doyon et al., 2008; Foley et al., 2009a; Foley et al., 2009b; Lawson and Wolfe, 2011; Moens et al., 2008; Sander et al., 2011a; Sander et al., 2011b; Sood et al., 2006). To gain access to this class of genes, we developed a Gateway (Kwan et al., 2007; Villefranc et al., 2007; Walhout et al., 2000) plasmid-based system for germline specific mutagenesis based on the system generated in the Zon lab (Ablain et al., 2015). CRISPR/Cas9 has become a standard method of mutagenesis in zebrafish and other organisms, and biallelic conversion events have been widely observed in mitotic cells (Ablain et al., 2015; Auer et al., 2014a; Auer et al., 2014b; Barrangou, 2013; Blackburn et al., 2013; Hruscha et al., 2013) et al., 2013; (Hwang et al., 2013a; Hwang et al., 2013b). However, the effectiveness of CRISPR/Cas9 in meiotic cells, when there are 4 copies of each chromosome and distinct checkpoints and repair pathways has not been determined.

Briefly, we generated mutagenesis cassettes that include selectable markers for the cassettes (tissue specific expression of fluorescent proteins (FP)) and that express target guide RNAs ubiquitously (under a U6 promoter) and Cas9 from germ cell specific promoters (*bucky ball* (meiotic cells in the female germline)(Heim et al., 2014) and *ziwi* (all germ cells)(Leu and Draper, 2010); Figure 1A,B, Supplemental Figure 1, 2). We developed two systems, one that marks only the germline and a second that includes the germline marker and an additional marker that labels the hearts of transgenic animals to allow for earlier selection (Figure 1A,C, Table 1). To generate transgenic animals, these cassettes along with *transposase* RNA were injected into 1-cell stage embryos to generate stable lines by Tol2 mediated integration of the transgene (Kawakami, 2005, 2007; Kawakami et al., 2004; Kawakami et al., 1998). We anticipated that mutations would be induced later by the *buc* mutagenesis cassette than with the *ziwi* mutagenesis cassette because the *buc* promoter is expressed only in females and in more advanced germ cells (Heim et al., 2014) than the *ziwi* promoter, which is expressed in mitotic germ cells (Leu and Draper, 2010). Hereafter, we refer to constructs and lines in which *cas9* is expressed from the *bucky ball* promoter as OMS for <u>o</u>vary <u>m</u>utagenesis <u>s</u>ystem and for those expressed from the *ziwi* promoter as GMS for <u>g</u>ermline <u>m</u>utagenesis <u>s</u>ystem. Because *buc* promoter is only activated in females, founder males can be used to propagate the transgenes and to generate additional mutant alleles in subsequent generations. This will be valuable for mutations that cause female sterile phenotypes (oocytes arrest and no eggs are produced) or if the maternal-effect phenotypes are not viable, like *buc* mutants. Significantly, even in cases where oocyte arrest is observed, histological assays can be used to pinpoint the precise stages that are affected because the transgenic oocytes (*oms*, *gms*) or sperm (*gms*) are marked with germline specific fluorescent reporters (Figure 1, 2). By sequencing the targeted region in marked oocytes or eggs, mutations induced in the germline can be identified.

**Figure 2.**
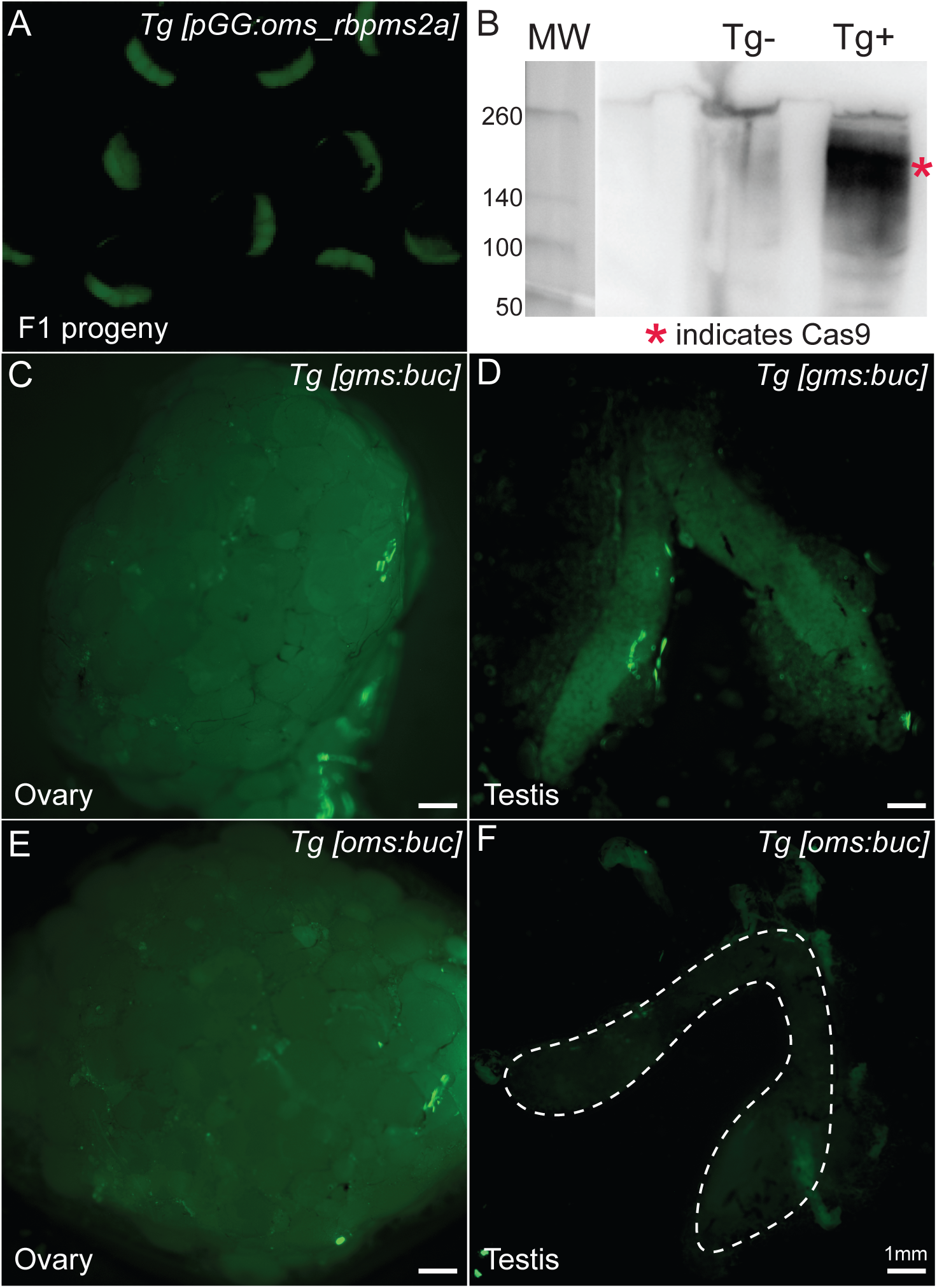
Ubiquitous Expression of Cas9 and Guide RNAs are not toxic to germ cell development or embryogenesis. A) Maternal expression of GFP in the F1 progeny of a transgenic mother. B) Western blot showing maternal expression of Cas9 protein in the progeny of a transgenic mother. C) Expression of GFP in ovary and D) testis of *gms* transgenic fish. E) Expression of GFP in the ovary but not the F) testis of *oms* transgenic fish.

### Validation and viability of OMS and GMS mutagenesis systems

In this manuscript we report cassettes and recovered transgenic (Tg+) founders targeting three genes, *rbpms2a, kif5Ba*, and *bucky ball* (Supplemental Table 1, 2) all of which have known germ cell functions. Founders were identified based on expression of the fluorescent reporter for the plasmid in their hearts and in the case of females, transmission of the germline reporter (GFP) to their progeny (Figure 2A, B). Although GFP was expected to be a proxy for Cas9 because both proteins are produced from the same transcript (Figure 1A), we confirmed that maternal Cas9 expression, like GFP, persists in embryos (Figure 2B). Based on our analysis of the GFP positive progeny of the founders and F1 that we have identified thus far, ubiquitous expression of the guide RNA is not toxic to germ cells (oocytes or sperm) or the resulting progeny nor is germline expression of Cas9 driven by *bucky ball (oms)* or *ziwi (gms)* promoters (Figure 2 A, Supplemental Figure 3).

### Phenocopy of *kif5Ba with* OMS and GMS vectors

We have previously shown that global loss of maternal *kif5Ba* (*Mkif5Ba)* results in failure to specify PGCs (Figure 3A) (Campbell et al., 2015). As a proof of concept and means to test the efficiency of our mutagenesis system, we used the vector mutagenesis systems to disrupt *kif5Ba*. We cloned a previously validated guide targeting the motor domain of *kif5Ba* (Campbell et al., 2015) into the *ziwi* promoter system II (hereafter called *Tg*:*GMS:kif5Ba*) (see Supplemental Tables 1, 2). We previously showed that the restriction enzyme MboI cuts the wild-type allele; however, when mutations are introduced in the targeted region, MboI can no longer cut the mutant allele (Campbell et al., 2015). Therefore, we used this assay to analyze the frequency of mutation in six Green heart positive F1 progeny of a *Tg:GMS:kif5Ba* male founder and their F2 progeny (Figure 3B, Supplemental Figure 3A,B). We isolated genomic DNA from somatic tissue (fin) and found that, as expected if no mutations were induced in the germline, five of the six F1 progeny were homozygous wild-type in their somatic tissues (Figure 3B). However, one of the females was heterozygous, indicating that *de novo* mutations in *kif5Ba* were made in her father’s sperm (Figure 3B). Because each of these F1 carry the *Tg:GMS:kif5Ba* mutagenesis transgene, we examined the progeny from pairwise intercrosses to screen for induced germline mutations (Figure 3B). Normally, a cross between two homozygous wild-type fish should yield only homozygous wild-type progeny while a cross between a homozygous wild-type fish and a heterozygote should yield half homozygous wild-type and half heterozygous progeny. Instead we found deviations from these expected genotypes in the progeny of intercrosses of *Tg:GMS:kif5Ba* carriers, indicative of CRISPR/Cas9 mediated mutation of the germ cells (Figure 3C). We next sequenced the F2 progeny to confirm that new mutations had been induced in sperm and oocytes of *Tg:GMS:kif5Ba* F1. Both in frame and deleterious mutations were recovered (Figure 3D).

**Figure 3.**
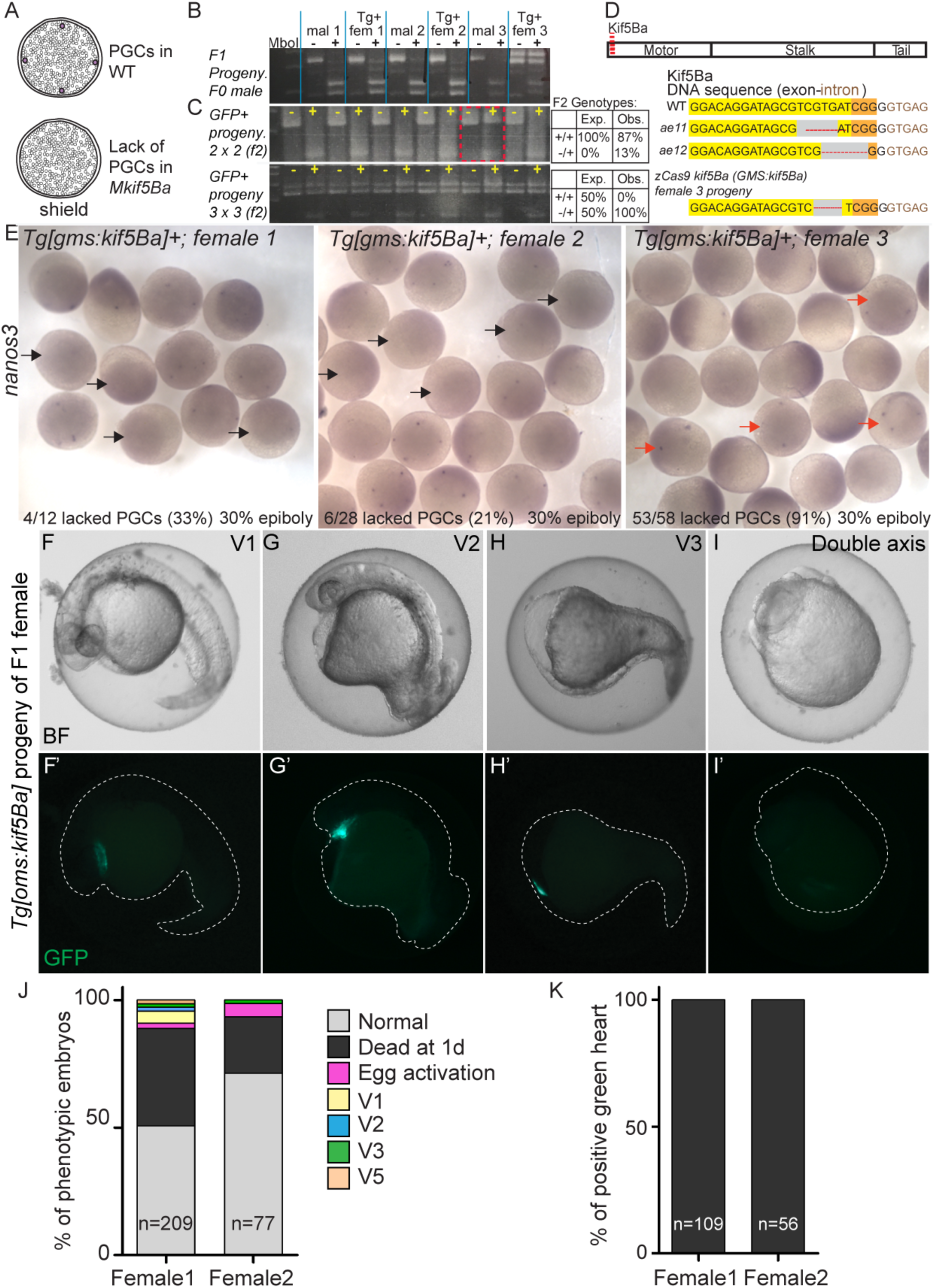
Phenocopy of maternal *kif5Ba* loss of function phenotypes. A) Schematic of wild-type and *Mkif5Ba* phenotypes. B,C) Representative restriction enzyme-based screens for mutations disrupting *kif5Ba* and expected and observed mutation frequencies in C) F1 and D) F2 progeny of transgenic parents. D) Schematic of Kif5Ba protein and targeted region and representative sequences from germline mutagenesis systems. D) *nanos3* staining in three independent clutches from F1 mothers carrying *Tg[ziwi:cas9T2Agfp_U6:kif5Baguide]* (*gms:kif5Ba).* Black arrows highlight progeny lacking germ cells in the first two panels and red arrows point out the few embryos with germ cells in the third panel. Progeny of F2 female transgenic for *Tg[ziwi:cas9T2Agfp_U6:kif5Baguide] (gms:kif5Ba)*. (F-F’) Embryos with a ventralization V1. (G-G’) Embryos with a ventralization V2. (H-H’) Embryos with a ventralization V3 (I-I’) Embryos with a ventralization V5. (J) Quantification of embryos ranging from normal to ventralization V5 stage. (K) Quantification of positive green heart of the progeny at d2.

Having confirmed that germline mutations were induced, we then examined the expression of the germ cell marker *nanos3* to determine if germ cells were present in the resulting maternal GFP positive progeny of three different *Tg:GMS:kif5Ba* F1 females (Figure 3). As expected for mosaic loss of maternal *kif5Ba* function, a fraction of the progeny from each *Tg:GMS:kif5Ba* F1 female lacked germ cells expressing *nanos3* (Figure 3E). The penetrance of phenotypic embryos was nonmendelian in number and varied from female to female (ranging from 21-91%), with the highest frequency of phenotypic progeny from the *Tg:GMS:kif5Ba* F1 mother that was already heterozygous for a mutation at the *kif5Ba* locus. In addition, maternal *kif5Ba* promotes dorso-ventral (DV) patterning by promoting the parallel vegetal microtubule array that mediates asymmetric distribution of the dorsal determinants toward the dorsal side (Campbell et al., 2015). To determine if *gms induced* alleles recapitulated this *Mkif5Ba* phenotype, we examined the phenotype of embryos from two different *Tg:GMS:kif5Ba* F1 females at d1. Dorso-ventral phenotypes ranging from mild to severe dorsalization and axis duplication (V1 to V5) were observed among the progeny of *Tg:GMS:kif5Ba* F1 females at d1 (Figure 3F-J), which were confirmed to carry the transgene based on their green hearts (Figure 3KF, Supplemental Figure 3). Based on these results, we conclude that *gms* induced mutations in *kif5Ba* can effectively phenocopy traditional maternal-effect loss of function *kif5Ba* phenotypes.

### Phenocopy of *buc*

Loss of *bucky ball* results in failure to establish the animal-vegetal axis of the oocyte and embryo (Bontems et al., 2009; Dosch et al., 2004; Heim et al., 2014; Marlow and Mullins, 2008). To test the OMS and GMS systems at another locus, we generated guide RNAs targeting exon 4 of the *bucky ball* gene and confirmed that the guides were mutagenic in the presence of Cas9 in transient embryo assays (Figure 4A, Supplemental Table 1, 2). Once mutagenic guides were identified, we cloned those guides into the OMS and GMS vectors and sequenced the resulting *oms:buc* and *gms:buc* plasmids (Figure 4A, Supplemental Table 1, 2). From 17 transient transgenics we recovered 1 female and 4 male founders for *gms:buc* (Figure 4B,C, Supplemental Figure 3C). The single female *gms:buc* founder produced embryos with either wild-type animal-vegetal polarity (n=22; 91.6%) or lacking an animal-vegetal axis (n=2; 8.4%) (Figure 4B, C, Supplemental Figure 3C). Thus, confirming that the *GMS* system can be used to generate and assess maternal-effect phenotypes in just one generation, a significant advantage over traditional screens for maternal-effect functions in which phenotypes are first detected after four generations (Dosch et al., 2004; Wagner et al., 2004) and haploid screens (Pelegri and Mullins, 2004; Pelegri and Schulte-Merker, 1999).

**Figure 4.**
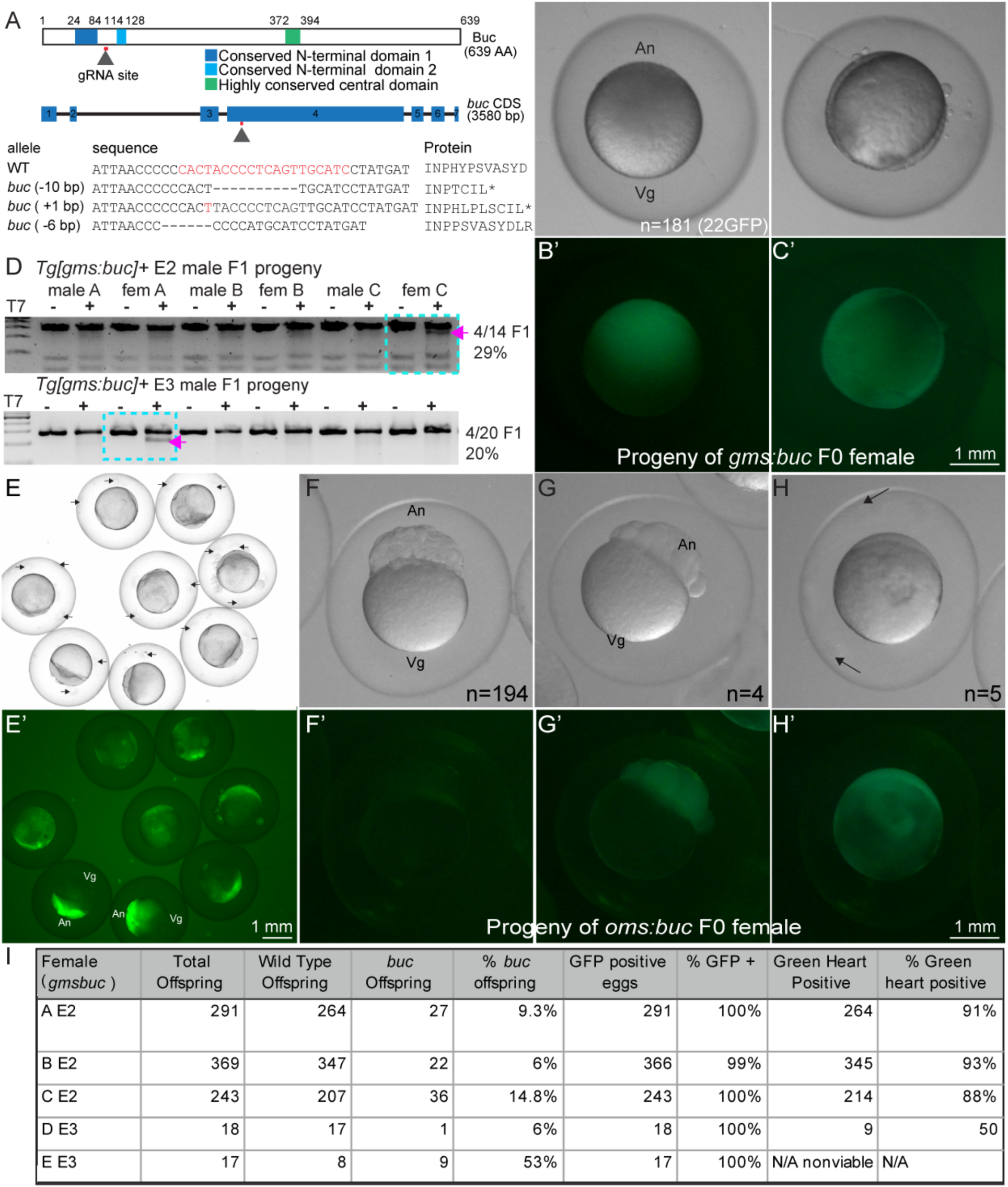
Phenocopy of loss of polarity due to mutation of *bucky ball* induced by GMS system. A) Schematic depicting site targeted in *bucky ball* and representative mutant alleles recovered. B,B’,C,C’) Progeny of F0 female transgenic for *Tg[ziwi:cas9T2Agfp_U6:bucguide]* (*gms:buc).* D) Representative T7 endonuclease assays to detect mutations in *bucky ball.* Mutagenesis frequencies are shown for F1 progeny of two independent founder males. E) Brightfield image and F) maternal GFP expression in a clutch from a F1 mother transgenic for *gms:buc.* F-H’) Progeny of F0 female transgenic for *Tg[buc:cas9T2Agfp_U6:bucguide]* (*oms:buc).* F-F’) Embryo with normal development and absence of transgene transmission (GFP negative). G-G’) Embryo with normal development and presence of transgene transmission (GFP positive). H-H’) Polar embryos with germline transmission of the transgene. (C-C’) Apolar embryos resulting from the germline expression and transmission of the transgene. I) Table showing transmission frequency of plasmids and penetrance of phenotypes in progeny of F1 mothers transgenic for *gms:buc.*

To detect de novo mutations T7 endonuclease assays were performed using genomic DNA from somatic tissues of the F1 progeny of two founder males. These analyses revealed 29% and 20% mutagenesis frequencies (Figure 4D), indicating that the expression of Cas9 from the ziwi promoter in the presence of guide RNA targeting *buc* induced mutations in the germline of the founder males. Sequencing of genomic DNA confirmed the mutations detected by the T7 assays and revealed that all four F1 progeny of founder male E2 (3 males and 1 female) carried the same mutation as one another and that four female offspring from founder male E3 with a mutant allele of *buc* carried a different mutation from E2, but the same mutation as one another (Figure 4D, Supplemental Figure 4). Next we examined the F1 females for *buc* phenotypes, specifically no animal-vegetal polarity and multiple micropyles, a somatic cell fate that is usually limited to one cell at the animal pole of wild-type but is ectopically specified in *buc* mutants (Heim et al., 2014; Marlow and Mullins, 2008). As expected for loss of *buc* function, five F1 females produced progeny with *bucky ball* phenotypes with penetrance ranging from 6-53% (Figure 4E, I).

Next we generated stable transgenic lines for the *oms:buc* alleles (Supplemental Figure 3 shows transgene transmission). We recovered several male founders carrying this transgene and one female founder who produced progeny with wild-type animal-vegetal axes (Figure 4F-G’) or lacking animal-vegetal polarity (Figure 4H, H’). As expected, based on activation of Cas9 at a later stage of germline development, the *oms* system, which is only activated in early meiotic oocytes, was less efficient than the *gms* system, which is activated in mitotic germ cells. Although the frequency of phenotype penetrance among individuals is somewhat variable for both *gms* and *oms* mutagenesis, ranging from lower than to greater than the penetrance of a typical recessive mutant phenotype, this approach represents a significant advance over germline replacement methods, which are tedious, time intensive, and inefficient.

## Overall Conclusions

Comparison of the OMS and GMS systems at two loci indicates that the GMS system, which drives expression in mitotic germ cells can achieve disruption of gene function specifically in the germline that yield phenotypic progeny with frequencies that match or exceed those of zygotic recessive alleles. In contrast, the OMS system, which drives expression in meiotic cells appears to be less efficient even with the same guide RNA. This is not surprising given that there are four copies of each chromosome in oocytes at this stage, and that all of them or the majority would need to be mutated to eliminate the maternal contribution. Nonetheless, this system may be suitable for genes that are first transcribed in late stage I or stage II oocytes. Significantly, detection of in-frame mutations indicates that phenotypic manifestation is an underrepresentation of the efficiency of mutagenesis, which encompasses both deleterious versus non-deleterious mutations. Nondisruptive mutations are potentially problematic in vector based loss of function studies because once the target site is altered it is no longer available for further mutagenesis. Here only a single guide RNA was used. One approach to overcome this potential limitation may be to generate a system in which there are multiple guides arrayed in tandem such that large regions of the gene can be deleted, thereby improving the frequency of generating loss of function germ cells and progeny lacking maternal gene function.

## Supporting information

Supplemental Figures,Legends, and Protocol

## Author Contributions

Experiments were conceived and designed by FLM and OHK. OHK generated the backbone vectors and *oms* and *gms kif5Ba* and *rbpms2* constructs and lines. SB designed *buc* guides and *oms:buc* and *gms:buc* lines. AS performed western blots for Cas9. SB, DD, KL, FLM performed all other experiments and analysis in consultation with FLM. FLM contributed reagents, materials, and analysis tools. All authors contributed to various aspects of data interpretation and discussion/editing of the manuscript. FLM and SB made the figures and wrote the manuscript.

## Acknowledgements

We thank members of the Marlow lab for helpful discussions, our animal care staff for fish care (Einstein and CCMS at ISMMS) and the Microscopy CoRE at Icahn School of Mount Sinai. This work was supported by National Institutes of Health, Grant R21-HD091456 to FLM. OHK was supported by T32-GM007288 and F30-HD082903.

## Notes

### Competing Interest Statement

The authors have declared no competing interest.

## References

Ablain, J., Durand, E.M., Yang, S., Zhou, Y., Zon, L.I., 2015. A CRISPR/Cas9 vector system for tissue-specific gene disruption in zebrafish. Developmental cell 32, 756–764.

Abrams, E.W., Fuentes, R., Marlow, F.L., Kobayashi, M., Zhang, H., Lu, S., Kapp, L., Joseph, S.R., Kugath, A., Gupta, T., Lemon, V., Runke, G., Amodeo, A.A., Vastenhouw, N.L., Mullins, M.C., 2020. Molecular genetics of maternally-controlled cell divisions. PLoS genetics 16, e1008652.

Ambartsumyan, G., Clark, A.T., 2008. Aneuploidy and early human embryo development. Human molecular genetics 17, R10–15.

Auer, T.O., Duroure, K., Concordet, J.P., Del Bene, F., 2014a. CRISPR/Cas9-mediated conversion of eGFP-into Gal4-transgenic lines in zebrafish. Nature protocols 9, 2823–2840.

Auer, T.O., Duroure, K., De Cian, A., Concordet, J.P., Del Bene, F., 2014b. Highly efficient CRISPR/Cas9-mediated knock-in in zebrafish by homology-independent DNA repair. Genome research 24, 142–153.

Barrangou, R., 2013. CRISPR-Cas systems and RNA-guided interference. Wiley interdisciplinary reviews. RNA 4, 267–278.

Bennett, J.T., Stickney, H.L., Choi, W.Y., Ciruna, B., Talbot, W.S., Schier, A.F., 2007. Maternal nodal and zebrafish embryogenesis. Nature 450, E1–2; discussion E2-4.

Blackburn, P.R., Campbell, J.M., Clark, K.J., Ekker, S.C., 2013. The CRISPR system--keeping zebrafish gene targeting fresh. Zebrafish 10, 116–118.

Bontems, F., Stein, A., Marlow, F., Lyautey, J., Gupta, T., Mullins, M.C., Dosch, R., 2009. Bucky ball organizes germ plasm assembly in zebrafish. Current biology: CB 19, 414–422.

Borovina, A., Superina, S., Voskas, D., Ciruna, B., Vangl2 directs the posterior tilting and asymmetric localization of motile primary cilia. Nature cell biology 12, 407–412.

Campbell, P.D., Heim, A.E., Smith, M.Z., Marlow, F.L., 2015. Kinesin-1 interacts with Bucky ball to form germ cells and is required to pattern the zebrafish body axis. Development 142, 2996–3008.

Cao, Z., Mao, X., Luo, L., 2019. Germline Stem Cells Drive Ovary Regeneration in Zebrafish. Cell reports 26, 1709–1717 e1703.

Ciruna, B., Jenny, A., Lee, D., Mlodzik, M., Schier, A.F., 2006. Planar cell polarity signalling couples cell division and morphogenesis during neurulation. Nature 439, 220–224.

Ciruna, B., Weidinger, G., Knaut, H., Thisse, B., Thisse, C., Raz, E., Schier, A.F., 2002. Production of maternal-zygotic mutant zebrafish by germ-line replacement. Proceedings of the National Academy of Sciences of the United States of America 99, 14919–14924.

Dosch, R., Wagner, D.S., Mintzer, K.A., Runke, G., Wiemelt, A.P., Mullins, M.C., 2004. Maternal control of vertebrate development before the midblastula transition: mutants from the zebrafish I. Developmental cell 6, 771–780.

Doyon, Y., McCammon, J.M., Miller, J.C., Faraji, F., Ngo, C., Katibah, G.E., Amora, R., Hocking, T.D., Zhang, L., Rebar, E.J., Gregory, P.D., Urnov, F.D., Amacher, S.L., 2008. Heritable targeted gene disruption in zebrafish using designed zinc-finger nucleases. Nature biotechnology 26, 702–708.

Dranow, D.B., Hu, K., Bird, A.M., Lawry, S.T., Adams, M.T., Sanchez, A., Amatruda, J.F., Draper, B.W., 2016. Bmp15 Is an Oocyte-Produced Signal Required for Maintenance of the Adult Female Sexual Phenotype in Zebrafish. PLoS genetics 12, e1006323.

Dranow, D.B., Tucker, R.P., Draper, B.W., 2013. Germ cells are required to maintain a stable sexual phenotype in adult zebrafish. Developmental biology 376, 43–50.

Foley, J.E., Maeder, M.L., Pearlberg, J., Joung, J.K., Peterson, R.T., Yeh, J.R., 2009a. Targeted mutagenesis in zebrafish using customized zinc-finger nucleases. Nature protocols 4, 1855–1867.

Foley, J.E., Yeh, J.R., Maeder, M.L., Reyon, D., Sander, J.D., Peterson, R.T., Joung, J.K., 2009b. Rapid mutation of endogenous zebrafish genes using zinc finger nucleases made by Oligomerized Pool ENgineering (OPEN). PloS one 4, e4348.

Hartung, O., Forbes, M.M., Marlow, F.L., 2014. Zebrafish vasa is required for germ-cell differentiation and maintenance. Molecular reproduction and development.

Hassold, T., Hall, H., Hunt, P., 2007. The origin of human aneuploidy: where we have been, where we are going. Human molecular genetics 16 Spec No. 2, R203–208.

Hassold, T., Hunt, P., 2007. Rescuing distal crossovers. Nature genetics 39, 1187–1188.

Heim, A.E., Hartung, O., Rothhamel, S., Ferreira, E., Jenny, A., Marlow, F.L., 2014. Oocyte polarity requires a Bucky ball-dependent feedback amplification loop. Development (Cambridge, England) 141, 842–854.

Hruscha, A., Krawitz, P., Rechenberg, A., Heinrich, V., Hecht, J., Haass, C., Schmid, B., 2013. Efficient CRISPR/Cas9 genome editing with low off-target effects in zebrafish. Development 140, 4982–4987.

Hwang, W.Y., Fu, Y., Reyon, D., Maeder, M.L., Kaini, P., Sander, J.D., Joung, J.K., Peterson, R.T., Yeh, J.R., 2013a. Heritable and precise zebrafish genome editing using a CRISPR-Cas system. PloS one 8, e68708.

Hwang, W.Y., Fu, Y., Reyon, D., Maeder, M.L., Tsai, S.Q., Sander, J.D., Peterson, R.T., Yeh, J.R., Joung, J.K., 2013b. Efficient genome editing in zebrafish using a CRISPR-Cas system. Nature biotechnology 31, 227–229.

Kaufman, O.H., Lee, K., Martin, M., Rothhamel, S., Marlow, F.L., 2018. rbpms2 functions in Balbiani body architecture and ovary fate. PLoS genetics 14, e1007489.

Kawakami, K., 2005. Transposon tools and methods in zebrafish. Developmental dynamics: an official publication of the American Association of Anatomists 234, 244–254.

Kawakami, K., 2007. Tol2: a versatile gene transfer vector in vertebrates. Genome biology 8 Suppl 1, S7.

Kawakami, K., Imanaka, K., Itoh, M., Taira, M., 2004. Excision of the Tol2 transposable element of the medaka fish Oryzias latipes in Xenopus laevis and Xenopus tropicalis. Gene 338, 93–98.

Kawakami, K., Koga, A., Hori, H., Shima, A., 1998. Excision of the tol2 transposable element of the medaka fish, Oryzias latipes, in zebrafish, Danio rerio. Gene 225, 17–22.

Kwan, K.M., Fujimoto, E., Grabher, C., Mangum, B.D., Hardy, M.E., Campbell, D.S., Parant, J.M., Yost, H.J., Kanki, J.P., Chien, C.B., 2007. The Tol2kit: a multisite gateway-based construction kit for Tol2 transposon transgenesis constructs. Developmental dynamics: an official publication of the American Association of Anatomists 236, 3088–3099.

Lawson, N.D., Wolfe, S.A., 2011. Forward and reverse genetic approaches for the analysis of vertebrate development in the zebrafish. Developmental cell 21, 48–64.

Leu, D.H., Draper, B.W., 2010. The ziwi promoter drives germline-specific gene expression in zebrafish. Developmental dynamics: an official publication of the American Association of Anatomists 239, 2714–2721.

Marlow, F.L., 2010. Maternal Control of Development in Vertebrates: My Mother Made Me Do It!, San Rafael (CA).

Marlow, F.L., Mullins, M.C., 2008. Bucky ball functions in Balbiani body assembly and animal-vegetal polarity in the oocyte and follicle cell layer in zebrafish. Developmental biology 321, 40–50.

Meeker, N.D., Hutchinson, S.A., Ho, L., Trede, N.S., 2007. Method for isolation of PCR-ready genomic DNA from zebrafish tissues. BioTechniques 43, 610, 612, 614.

Moens, C.B., Donn, T.M., Wolf-Saxon, E.R.,, Ma, T.P., 2008. Reverse genetics in zebrafish by TILLING. Brief Funct Genomic Proteomic 7, 454–459.

Pelegri, F., Dekens, M.P., Schulte-Merker, S., Maischein, H.M., Weiler, C., Nusslein-Volhard, C., 2004. Identification of recessive maternal-effect mutations in the zebrafish using a gynogenesis-based method. Developmental dynamics: an official publication of the American Association of Anatomists 231, 324–335.

Pelegri, F., Mullins, M.C., 2004. Genetic screens for maternal-effect mutations. Methods in cell biology 77, 21–51.

Pelegri, F., Schulte-Merker, S., 1999. A gynogenesis-based screen for maternal-effect genes in the zebrafish, Danio rerio. Methods in cell biology 60, 1–20.

Rodriguez-Mari, A., Canestro, C., Bremiller, R.A., Nguyen-Johnson, A., Asakawa, K., Kawakami, K., Postlethwait, J.H., 2010. Sex reversal in zebrafish fancl mutants is caused by Tp53-mediated germ cell apoptosis. PLoS genetics 6, e1001034.

Rodriguez-Mari, A., Postlethwait, J.H., 2011. The role of Fanconi anemia/BRCA genes in zebrafish sex determination. Methods in cell biology 105, 461–490.

Rodriguez-Mari, A., Wilson, C., Titus, T.A., Canestro, C., BreMiller, R.A., Yan, Y.L., Nanda, I., Johnston, A., Kanki, J.P., Gray, E.M., He, X., Spitsbergen, J., Schindler, D., Postlethwait, J.H., 2011. Roles of brca2 (fancd1) in oocyte nuclear architecture, gametogenesis, gonad tumors, and genome stability in zebrafish. PLoS genetics 7, e1001357.

Sander, J.D., Cade, L., Khayter, C., Reyon, D., Peterson, R.T., Joung, J.K., Yeh, J.R., 2011a. Targeted gene disruption in somatic zebrafish cells using engineered TALENs. Nature biotechnology 29, 697–698.

Sander, J.D., Yeh, J.R., Peterson, R.T., Joung, J.K., 2011b. Engineering zinc finger nucleases for targeted mutagenesis of zebrafish. Methods in cell biology 104, 51–58.

Sato, Y., Hilbert, L., Oda, H., Wan, Y., Heddleston, J.M., Chew, T.L., Zaburdaev, V., Keller, P., Lionnet, T., Vastenhouw, N., Kimura, H., 2019. Histone H3K27 acetylation precedes active transcription during zebrafish zygotic genome activation as revealed by live-cell analysis. Development 146.

Siegfried, K.R., Nusslein-Volhard, C., 2008. Germ line control of female sex determination in zebrafish. Developmental biology 324, 277–287.

Slanchev, K., Stebler, J., de la Cueva-Mendez, G., Raz, E., 2005. Development without germ cells: the role of the germ line in zebrafish sex differentiation. Proceedings of the National Academy of Sciences of the United States of America 102, 4074–4079.

Sood, R., English, M.A., Jones, M., Mullikin, J., Wang, D.M., Anderson, M., Wu, D., Chandrasekharappa, S.C., Yu, J., Zhang, J., Paul Liu, P., 2006. Methods for reverse genetic screening in zebrafish by resequencing and TILLING. Methods 39, 220–227.

Thisse, B., Heyer, V., Lux, A., Alunni, V., Degrave, A., Seiliez, I., Kirchner, J., Parkhill, J.P., Thisse, C., 2004. Spatial and temporal expression of the zebrafish genome by large-scale in situ hybridization screening. Methods in cell biology 77, 505–519.

Vastenhouw, N.L., Cao, W.X., Lipshitz, H.D., 2019. The maternal-to-zygotic transition revisited. Development 146.

Villefranc, J.A., Amigo, J., Lawson, N.D., 2007. Gateway compatible vectors for analysis of gene function in the zebrafish. Developmental dynamics: an official publication of the American Association of Anatomists 236, 3077–3087.

Wagner, D.S., Dosch, R., Mintzer, K.A., Wiemelt, A.P., Mullins, M.C., 2004. Maternal control of development at the midblastula transition and beyond: mutants from the zebrafish II. Developmental cell 6, 781–790.

Walhout, A.J., Temple, G.F., Brasch, M.A., Hartley, J.L., Lorson, M.A., van den Heuvel, S., Vidal, M., 2000. GATEWAY recombinational cloning: application to the cloning of large numbers of open reading frames or ORFeomes. Methods in enzymology 328, 575–592.

Weidinger, G., Stebler, J., Slanchev, K., Dumstrei, K., Wise, C., Lovell-Badge, R., Thisse, C., Thisse, B., Raz, E., 2003. dead end, a novel vertebrate germ plasm component, is required for zebrafish primordial germ cell migration and survival. Current biology: CB 13, 1429–1434.

Williams, M.L.K., Sawada, A., Budine, T., Yin, C., Gontarz, P., Solnica-Krezel, L., 2018. Gon4l regulates notochord boundary formation and cell polarity underlying axis extension by repressing adhesion genes. Nature communications 9, 1319.

Wu, D., Dean, J., 2020. Maternal factors regulating preimplantation development in mice. Current topics in developmental biology 140, 317–340.

Wu, E., Vastenhouw, N.L., 2020. From mother to embryo: A molecular perspective on zygotic genome activation. Current topics in developmental biology 140, 209–254.

Zhao, L.Y., Niu, Y., Santiago, A., Liu, J., Albert, S.H., Robertson, K.D., Liao, D., 2006. An EBF3-mediated transcriptional program that induces cell cycle arrest and apoptosis. Cancer research 66, 9445–9452.

