## Supplemental Figures,Legends, and Protocol for "A Transgenic System for Targeted Ablation of Reproductive and Maternal-Effect genes"

**Bertho and Kaufman et al -Supplemental Figures and legends**

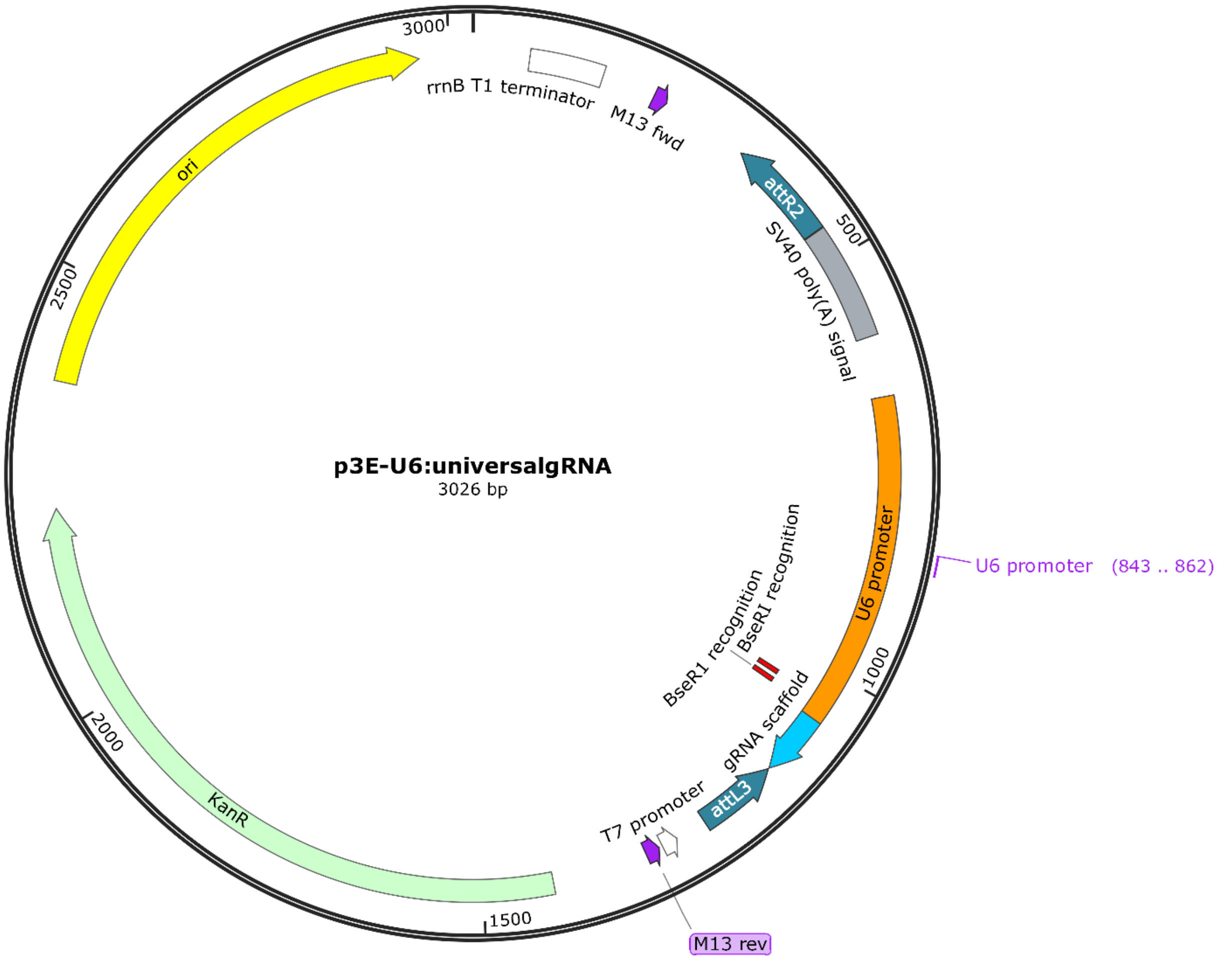


**Supplemental Figure 1. Plasmid map of p3E_polyA_U6:universalgRNA.** The plasmid contains attR2 and attL3 recombination sites (dark turquoise), SV40 poly(A) signal (grey), U6 promoter (orange), BseR1 recognition site (red) where the specific gRNA will be placed and the gRNA scaffold (light blue) and the sequencing primer (Forward direction).

**Supplemental Figure 2. P3E_polyA_U6:universalgRNA sequence.**

attR2 and attL3 (green), SV 40 polyA (grey), U6 promoter (orange), primer (purple), BseR1 enzyme site (red), gRNA scaffold (blue).

TGGCGGGCGTCCTGCCCGCCACCCTCCGGGCCGTTGCTTCACAACGTTCAAATCCGCTCCCGGCG
GATTTGTCCTACTCAGGAGAGCGTTCACCGACAAACAACAGATAAAACGAAAGGCCCAGTCTTCCGA
CTGAGCCTTTCGTTTTATTTGATGCCTGGCAGTTCCCTACTCTCGCGTTAACGCTAGCATGGATGTTT
TCCCAGTCACGACGTTGTAAAACGACGGCCAGTCTTAAGCTCGGGCCCTGCAGCTCTAGAGCTCGA
ATTCTACAGGTCACTAATACCATCTAAGTAGTTGGTTTACAGGTCACTAATACCATCTAAGTAGTTGGT
TCATAGTGACTGCATATGTTGTGTTTTACAGTATTATGTAGTCTGTTTTTTATGCAAAATCTAATTTAAT
ATATTGATATTTATATCATTTTACGTTTCTCGTTCAACTTTCTTGTACAAAGTGGGGGATCCAGACATG
ATAAGATACATTGATGAGTTTGGACAAACCACAACTAGAATGCAGTGAAAAAAATGCTTTATTTGTGA
AATTTGTGATGCTATTGCTTTATTTGTAACCATTATAAGCTGCAATAAACAAGTTAACAACAACAATTG
CATTCATTTTATGTTTCAGGTTCAGGGGGAGGTGTGGGAGGTTTTTTCATCATCGATGCGTCTTTTGT
TCTGGTCATCAAGGAGGGGGAATGTTCTGCGCATGCCTGTGGGGGAGGGAGAAGGACACGTCAC
TGAAAACGTCCCTGCATCACACCGAGACACCCAATCACTCAAGCCGAGACCAGATAATTTTGCATAT
GCTTTACAGTTTGAAAAATACCACGGTAAACCCTCACACAAACTCTGGATTTGAGATCTTTCAGGTTT
TATCAGTTTGCAGGTTTATGTCACCATGATATAGGGTCAGACTTGATCTAAGGGAGCTGAATAAGTGG
TTTAGTCACTCACCACCTCCCAAAAACATACCCAGAAGTCCCTGGTATATATAGCTCTCCCTCCAGCT
CTTGGTTCGGCTAGCACTCCTCAGTGACGAGGAGTCAGTAGCGTTTTAGAGCTAGAAATAGCAAGTT
AAAATAAGGCTAGTCCGTTATCAACTTGAAAAAGTGGCACCGAGTCGGTGCTCAACTTTATTATACAA
AGTTGGCATTATAAAAAAGCATTGCTTATCAATTTGTTGCAACGAACAGGTCACTATCAGTCAAAATAA
AATCATTATTTGGAGCTCCATGGTAGCGTTAACGCGGCCGCGATATCCCCTATAGTGAGTCGTATTA
CATGGTCATAGCTGTTTCCTGGCAGCTCTGGCCCGTGTCTCAAAATCTCTGATGTTACATTGCACAA
GATAAAAATATATCATCATGAACAATAAAACTGTCTGCTTACATAAACAGTAATACAAGGGGTGTTATG
AGCCATATTCAACGGGAAACGTCGAGGCCGCGATTAAATTCCAACATGGATGCTGATTTATATGGGT
ATAAATGGGCTCGCGATAATGTCGGGCAATCAGGTGCGACAATCTATCGCTTGTATGGGAAGCCCG
ATGCGCCAGAGTTGTTTCTGAAACATGGCAAAGGTAGCGTTGCCAATGATGTTACAGATGAGATGGT
CAGACTAAACTGGCTGACGGAATTTATGCCTCTTCCGACCATCAAGCATTTTATCCGTACTCCTGATG
ATGCATGGTTACTCACCACTGCGATCCCCGGAAAAACAGCATTCCAGGTATTAGAAGAATATCCTGA
TTCAGGTGAAAATATTGTTGATGCGCTGGCAGTGTTCCTGCGCCGGTTGCATTCGATTCCTGTTTGTA
ATTGTCCTTTTAACAGCGATCGCGTATTTCGTCTCGCTCAGGCGCAATCACGAATGAATAACGGTTTG
GTTGATGCGAGTGATTTTGATGACGAGCGTAATGGCTGGCCTGTTGAACAAGTCTGGAAAGAAATGC
ATAAACTTTTGCCATTCTCACCGGATTCAGTCGTCACTCATGGTGATTTCTCACTTGATAACCTTATTT
TTGACGAGGGGAAATTAATAGGTTGTATTGATGTTGGACGAGTCGGAATCGCAGACCGATACCAGGA

TCTTGCCATCCTATGGAACTGCCTCGGTGAGTTTTCTCCTTCATTACAGAAACGGCTTTTTCAAAAAT
ATGGTATTGATAATCCTGATATGAATAAATTGCAGTTTCATTTGATGCTCGATGAGTTTTTCTAATCAG
AATTGGTTAATTGGTTGTAACACTGGCAGAGCATTACGCTGACTTGACGGGACGGCGCAAGCTCATG
ACCAAAATCCCTTAACGTGAGTTACGCGTCGTTCCACTGAGCGTCAGACCCCGTAGAAAAGATCAAA
GGATCTTCTTGAGATCCTTTTTTTCTGCGCGTAATCTGCTGCTTGCAAACAAAAAAACCACCGCTACC
AGCGGTGGTTTGTTTGCCGGATCAAGAGCTACCAACTCTTTTTCCGAAGGTAACTGGCTTCAGCAGA
GCGCAGATACCAAATACTGTCCTTCTAGTGTAGCCGTAGTTAGGCCACCACTTCAAGAACTCTGTAG
CACCGCCTACATACCTCGCTCTGCTAATCCTGTTACCAGTGGCTGCTGCCAGTGGCGATAAGTCGTG
TCTTACCGGGTTGGACTCAAGACGATAGTTACCGGATAAGGCGCAGCGGTCGGGCTGAACGGGGG
GTTCGTGCACACAGCCCAGCTTGGAGCGAACGACCTACACCGAACTGAGATACCTACAGCGTGAGC
ATTGAGAAAGCGCCACGCTTCCCGAAGGGAGAAAGGCGGACAGGTATCCGGTAAGCGGCAGGGTC
GGAACAGGAGAGCGCACGAGGGAGCTTCCAGGGGGAAACGCCTGGTATCTTTATAGTCCTGTCGG
GTTTCGCCACCTCTGACTTGAGCGTCGATTTTTGTGATGCTCGTCAGGGGGGCGGAGCCTATGGAA
AAACGCCAGCAACGCGGCCTTTTTACGGTTCCTGGCCTTTTGCTGGCCTTTTGCTCACATGTT

A F0 transmission frequencies

| Founder - system I | Transmission freq - GH positive |
|---|---|
| Tg[pGG:buc:cas_U6rbpms2a gRNA] 2681-6 female | 0.53 |
| Tg[pGG:buc:cas_U6rbpms2a gRNA] 2681-2 male | 0.13 |
| Tg[pGG:buc:cas_U6rbpms2a gRNA] 2682-5 male | 0.25 |
| Tg[pGG:buc:cas_U6rbpms2a gRNA] 2682-9 male | 0.04 |
| Tg[pGG:ziwi:cas_U6rbpms2a gRNA] 2706-14 | tbc |
| Tg[pGG:ziwi:cas_U6rbpms2a gRNA] 2705-6 | 0.25 |
| Tg[pGG:ziwi:cas_U6rbpms2a gRNA] 2705-7 | 0.25 |

| Founder - system II | Transmission freq - GH positive |
|---|---|
| Tg[pGH:Ziwi:cas9T2AGFP_U6kif5Ba gRNA] 3073-1 male | 0.27 |
| Tg[pGH:Ziwi:cas9T2AGFP_U6kif5Ba gRNA] 3590-1 male | 0.01 |
| Tg[pGH:Ziwi:cas9T2AGFP_U6kif5Ba gRNA] 3590-5-20 male | 0.03 |
| Tg[pGH:Ziwi:cas9T2AGFP_U6rbpms2a gRNA] 3074-4 female | 0.29 |
| Tg[pGH:Ziwi:cas9T2AGFP_U6rbpms2a gRNA] 3074-7 female | ND |

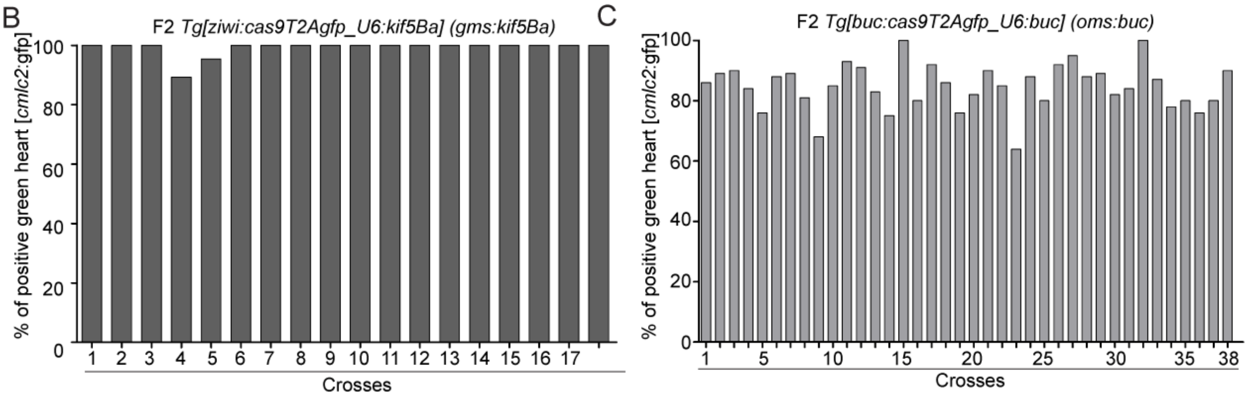


**Supplemental Figure 3. Germline transgene transmission.** A) Representative F0 transmission frequencies (GFP positive hearts). B) Quantification of positive green heart expression among*Tg[buc:cas9T2Agfp_U6:buc]* (*oms:buc)* embryos (n=18). C) Quantification of positive green heart expression among *Tg[ziwi:cas9T2Agfp_U6:kif5Ba] (gms:kif5Ba)* embryos (n=38).

A

| allele | sequence | Protein |
|---|---|---|
| WT | ATTAACCCCCACTACCCCTCAGTTGCATCCTATGAT | INPHYPSVASYD |
| *buc* (-10 bp) | ATTAACCCCCACT----------TGCATCCTATGAT | INPTCIL* |
| *buc* ( +1 bp) | ATTAACCCCCACTTACCCCTCAGTTGCATCCTATGAT | INPHLPLSCIL* |
| *buc* ( -6 bp) | ATTAACCC------CCCCATGCATCCTATGAT | INPPSVASYDLR |

B

WT - Chromatogram

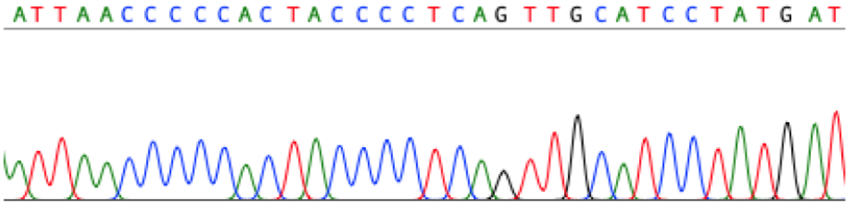


*buc* ( - 10 bp) - Chromatogram

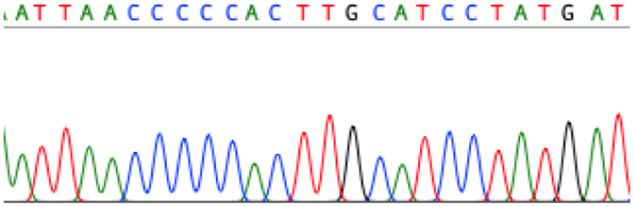


*buc* ( +1 bp) - Chromatogram

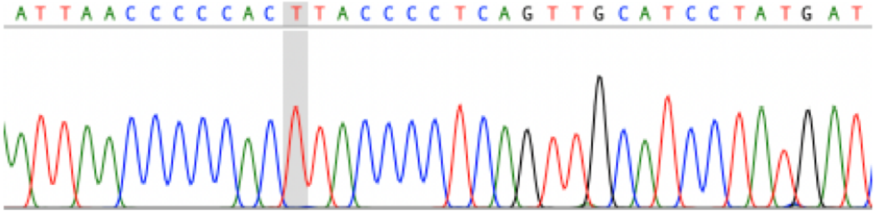


*buc* ( - 6 bp) - Chromatogram

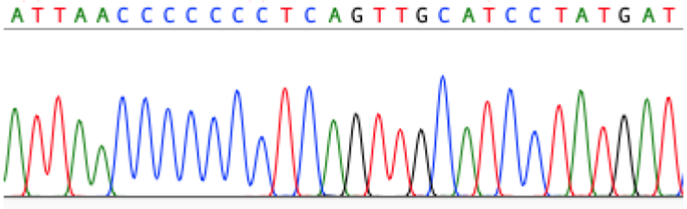


**Supplemental Figure 4. Sequence and chromatogram from recovered *buc* germline mutations from *gms:buc* mothers.** (A) Sequence of the region flanking the *buc* gRNA site and recovered sequences after Cas9 cleavage. (B) Chromatogram of the WT sequence and the mutated sequences in eggs from *gms:buc* mothers.

**Supplemental Tables:**

**Supplemental Table 1. Plasmids generated/used in this study**

| Marlow Lab Database Number | Plasmid Name | Other Info. |
|---|---|---|
| #60 | *p5E pBD119 ziwi promoter* | (Leu and Draper, 2010) |
| #190 | *2K Buc promoter in PDONR (p5E-Buc)* | (Heim et al., 2014) |
| #878 | *Tol2-R4/R3 cmcl2:gfp* | Tol2 sites surrounding R4/R3 att sites, with cmcl2:gfp transgenesis marker, Tol2 v1.0 (Kwan et al., 2007) |
| #1190 | *pME-Cas9* | Addgene 63154 (Zon lab) (Ablain et al., 2015) |
| #1191 | *pME-cas9-T2A-GFP* | Addgene 63155 (Zon lab) (Ablain et al., 2015) |
| #1194 | *p3E_pA_U6:rbpms2a_gRNA* | Guide from (Kaufman et al., 2018) |
| #1195 | *p3E_pA_U6:kif5Ba_gRNA* | Guide from (Campbell et al., 2015) |
| #1201 | *pTol_ZG_ziwi:cas9_u6:rbpms2a_gRNA* | Guide from (Kaufman et al., 2018) |
| #1202 | *pTol_ZG_buc:cas9_u6:rbpms2a_gRNA* | Guide from (Kaufman et al., 2018) |
| #1208 | *pGH_ubi_cas9 u6 :rbpms2a gRNA #5* | |
| #1244 | *pGH ziwi:cas9-T2A-GFP u6:kif5Ba gRNA (GMS)* | Guide from (Campbell et al., 2015) |
| #1245 | *pGH buc:cas9-T2A-GFP u6:kif5Ba gRNA (OMS)* | Guide from (Campbell et al., 2015) |

| Marlow Lab Database Number | Plasmid Name | Other Info. |
|---|---|---|
| #1246 | *p3E_pA_u6:universal gRNA* | |
| #1268 | *pGHziwiCas9T2AGFPU6bucgRNA* | |
| #1269 | *pGHbucCas9T2AGFPU6bucgRNA* | |
| #1272 | *p3E-pA-U6bucgRNA* | |

### Supplemental Table 2. Primers and guide RNAs used in this study

| **Marlow Lab Number** | **Primer Name** | **Primer Sequence (5' to 3')** |
|---|---|---|
| B9C4 | *bucprmtrinternalF* | CGGTGTTTTGTATCCTGTGC |
| B9C6 | *ziwiprminternalF* | CGTGAGGGTGAGTAAATAGTGTG |
| B17B5 | *pTol_u6gRNA_seqP* | CCTCACACAAACTCTGGATT |
| B20A10 | *cas9int_R1* | AATCTGATTGGAGCCCTGCT |
| - | *M13R (used to verify gRNA ligation into #1246)* | GCGGATAACAATTTCACACAGG |
| B28F4 | *bucex4crisprf2f* | TGCAGTATCCTGGCTATGTGAT |
| B28F5 | *bucex4crisprf2r* | ACCACATCAGGGGTAGAAGAGA |
| B10F7 | *kif5baex1CR-d-F1* | GGAGTGCACCATTAAAGTCATGTG |
| B10F10 | *kif5baex1CR-d-R2F2* | GTGTCGGTGTCAAATATTGAGGTC |
| B20G1 | *gRNA-bucex4-GMS-Fwd1* | GATGCAACTGAGGGGTAGTGGT |
| B20G2 | *gRNA-bucex4-GMS-Rev1* | CACTACCCCTCAGTTGCATCGA |
| B16A9 | *crispr constant (Gagnon et al., 2014a)* | AAAAGCACCGACTCGGTGCCACTTTTTCAAGTTGATAACGGACTAGCCTTATTTTAACTTGCTATTTCTAGCTCTAAAAC |
| B19F3 | *bucex4crispr1* | ATTTAGGTGACACTATACCACCCAGTAAACCACACAAGACGTTTTAGAGCTAGAAATAGCAAG |
| B19F4 | *bucex4crispr2* | ATTTAGGTGACACTATACCCCCACTACCCCTCAGTTGCATGTTTTAGAGCTAGAAATAGCAAG |
| B20B9 | *bucex4crispr3* | ATTTAGGTGACACTATAGGAATAAGCACTGCCAATTGTTTTAGAGCTAGAAATAGCAAG |
| B20B10 | *bucex4crispr4* | ATTTAGGTGACACTATAGGAGAACATTTCATCTCTGGTTTTAGAGCTAGAAATAGCAAG |

| Marlow Lab Number | gRNA Name | gRNA Sequence |
| --- | --- | --- |
| B16A9 | *crispr constant* (Gagnon et al., 2014a) | AAAAGCACCGACTCGGTGCCACTTTTTCAAG TTGATAACGGACTAGCCTTATTTTAACTTGCT ATTTCTAGCTCTAAAAC |
| B19F3 | *bucex4crispr1* | ATTTAGGTGACACTATACCACCCAGTAAACCA CACAAGACGTTTTAGAGCTAGAAATAGCAAG |
| B19F4 | *bucex4crispr2* | ATTTAGGTGACACTATACCCCCACTACCCCTC AGTTGCATGTTTTAGAGCTAGAAATAGCAAG |
| B20B9 | *bucex4crispr3* | ATTTAGGTGACACTATAGGAATAAGCACTGC CAATTGTTTTAGAGCTAGAAATAGCAAG |
| B20B10 | *bucex4crispr4* | ATTTAGGTGACACTATAGGAGAACATTTCATC TCTGGTTTTAGAGCTAGAAATAGCAAG |

**Supplemental Methods:**

### Protocol for germline-specific gene disruption in zebrafish

**Adapted from (Ablain et al., 2015)**

#### Overview

This protocol is based on (Ablain et al., 2015) tissue-specific gene disruption protocol in zebrafish. The method allows gene inactivation in zebrafish in a germline-specific manner. It can be used to analyze maternal effect genes in F1 embryos and generate stable tissue-specific knock-out lines analyze maternal effect genes. It takes advantage of the Tol2 transposase technology to integrate in the fish genome a vector expressing a guide RNA (gRNA) from a ubiquitous zebrafish U6 promoter and Cas9 under the control of a tissue-specific promoter. This protocol comprises 5 steps: 1) the identification of efficient CRISPR target sequences in the gene of interest; 2) the annealing of gene-specific oligonucleotides; 3) the construction of the germline-specific CRISPR vector; 4) the injection of the CRISPR construct in zebrafish embryos; 5) the phenotypic analysis and generation of stable lines (Fig. P1).

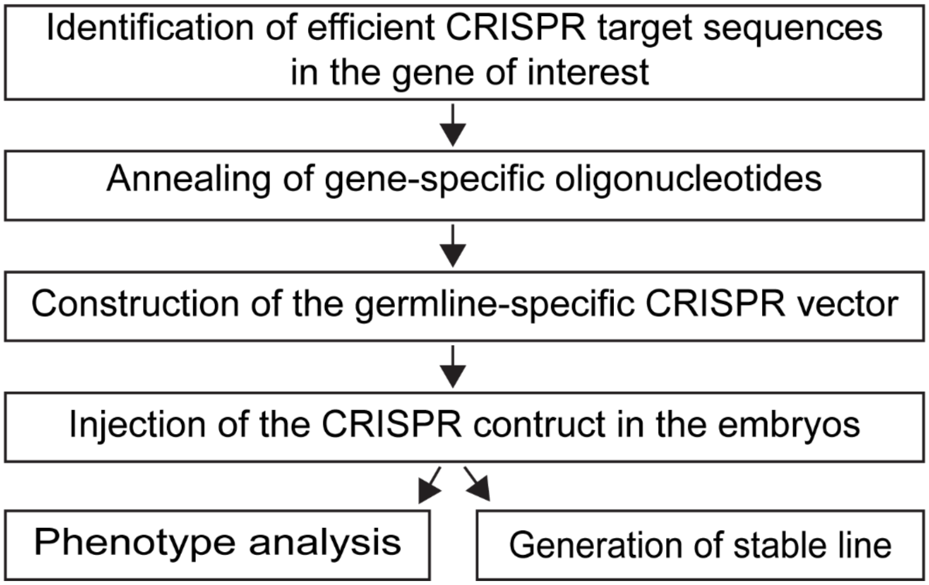


**Figure P1 – Workflow of germline-specific gene disruption**

**Reagents**

- Destination vector for Gateway (e.g. pDestTol2CG2)

- Middle entry vector for Gateway containing Cas9 (e.g. pME-Cas9 or pME-Cas9-T2A-GFP, available through Addgene)

- 5' entry vector for Gateway containing the germline-specific promoter of interest (p5E_*ziwi* promoter or p5E_*buc* promoter)

- 3' entry vector for Gateway containing a polyA sequence (p3E_polyA_U6:universalgRNA)

- Gene-specific oligonucleotides (see below)

- BseRI enzyme (New England Biolabs)

- Gateway LR clonase II (Invitrogen)

- T4 DNA ligase (New England Biolabs)

- Gel extraction kit or PCR-purification kit (Qiagen)

**Procedure**

All the steps except step 3 are identical to Ablain et al procedure. (Ablain et al., 2015)

1. Identification of efficient CRISPR target sequences in the gene(s) of interest

Pick several (3 to 6) CRISPR target sequences in each gene of interest using available tools (Hsu et al., 2013; Montague et al., 2014). Produce the gRNAs *in vitro* as per usual procedures (Gagnon et al., 2014b; Hwang et al., 2013). Inject the gRNAs along with Cas9 protein or mRNA into one-cell stage embryos of a WT strain. Extract DNA from injected embryos at 24 or 48 hpf by the HotSHOT method (Meeker et al., 2007) and assess mutation rates at target loci by sequencing or enzymatic assays (e.g. T7E1 assay, Surveyor assay) (Gagnon et al., 2014b; Kim et al., 2009). Proceed with the target sequences that have shown effective targeting (we usually only keep target sequences for which the mutation rates exceed 10%).

***Note:*** *alternatively, it is possible to test the efficiency of target sequences directly in the context of the Tol2 vector by cloning various target sequences in the U6:gRNA cassette of a vector expressing Cas9 under the control of a ubiquitous promoter.*

2. Annealing of gene-specific oligonucleotides

Order 22-mer, unmodified oligonucleotides as follows:
Forward: target sequence (20 bases)-GT
Reverse: reverse complement of target sequence-GA
***Note:*** *the target sequence must start with a G. If not, replace the first base by a G. This will introduce a mismatch but most mismatches at the 5' end of the target sequence are well tolerated.*

Example: for CRISPR target sequence GGTGGGAGAGTGGATGGCTG, order GGTGGGAGAGTGGATGGCTGGT (forward) and CAGCCATCCACTCTCCCACCGA (reverse).
Anneal the two oligos in thermocycler (5 min. at 95°C, -1°C/min. down to 20°C).

3. Construction of the germline-specific CRISPR vector

Clone the gene-specific seed sequence into the p3E_polyA_U6:universal gRNA predigested with BseRI and then perform the Gateway reaction with the germline-specific promoter (ziwi or buc) and Cas9 (Fig. P2).

- Digest the p3E_polyA_U6:universalgRNA of your choice with BseRI enzyme.
- Purify on 1% agarose gel or column.
- Ligate the gene-specific seed sequence into BseRI-digested p3E_polyA_U6:universalgRNA vector (1:20 vector:insert molar ratio).
- Transform chemically competent bacteria, plate on LB Agar and select with Kanamycin

- Extract plasmid DNA from 2-3 distinct colonies and sequence using the following primer: CCTCACACAAACTCTGGATT to check the insertion of the specific gRNA.
- Perform the Gateway reaction (Hartley et al., 2000) with the destination vector, a 5' entry vector containing a tissue-specific promoter of interest, a middle entry vector containing zebrafish codon-optimized Cas9 and p3E-polyA, according to manufacturer's protocol.
- Transform chemically competent bacteria, plate on LB Agar and select with ampicillin.
- Extract plasmid DNA from 2-3 colonies and check correct recombination by digestion or sequencing (*ziwi* or *buc*, *Cas9*, *U6* promoter primer listed in the table).

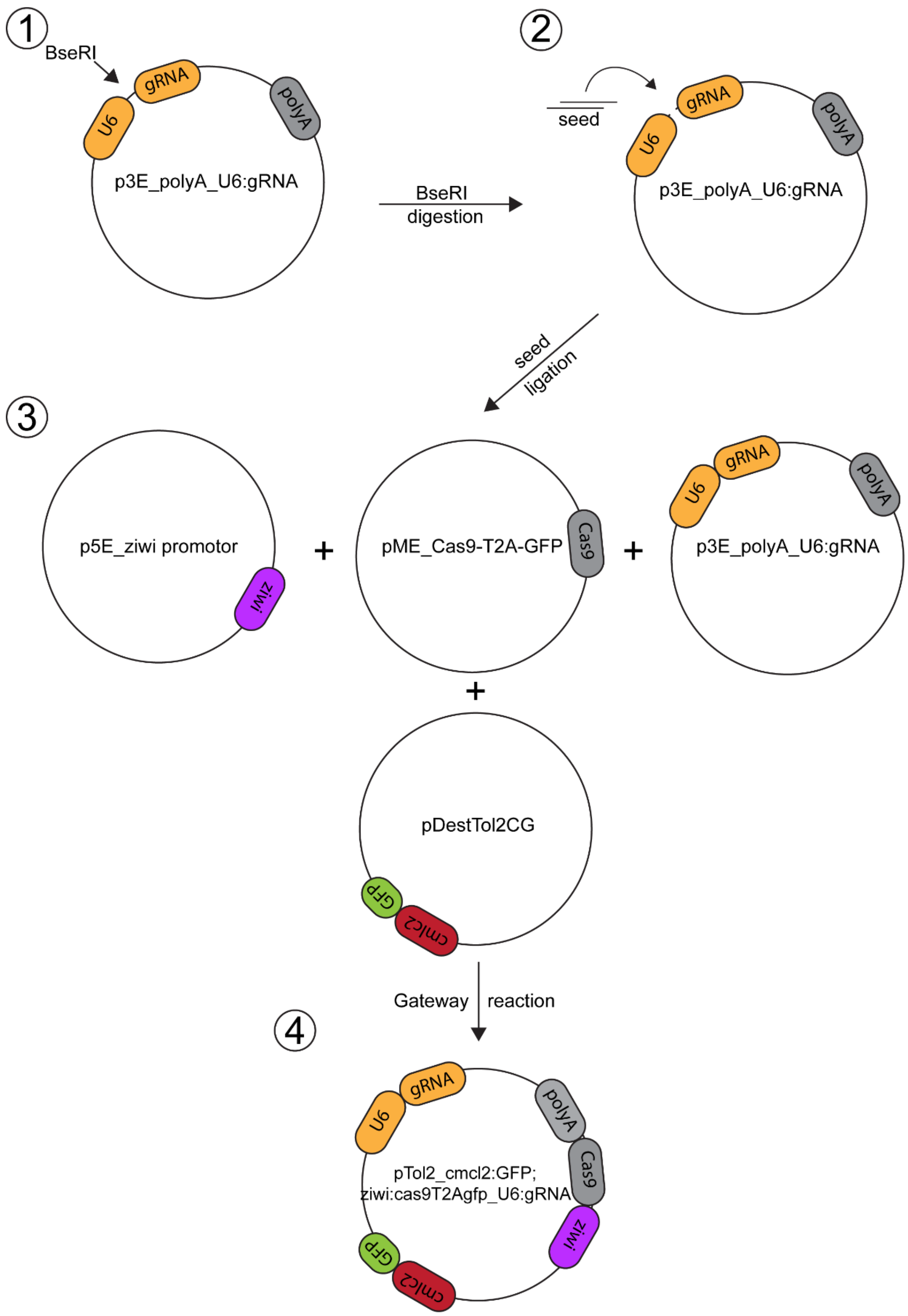


**Figure P2: Construction of a germline-specific CRISPR vector**

4. Injection of the CRISPR construct in zebrafish embryos

Mix and inject 20-30 pg of CRISPR vector and 20 pg of Tol2 mRNA into one-cell stage embryos (Kawakami et al., 2004). For reliable phenotypic analyses, we recommend injecting >50 embryos per construct. Vectors expressing gRNAs targeting an irrelevant gene or driving Cas9 expression in a different tissue can be used as negative controls. Allow injected embryos to develop at 28.5°C.

5. Phenotypic analysis and generation of stable lines

Sort injected F0 embryos based on the expression of a transgenesis marker present either in the destination vector (e.g. cmlc2:GFP) or in the middle entry part of the vector (e.g. T2A-GFP). Only positive embryos should be considered for further analysis.
For the generation of stable lines, raise positive F0 fish to adulthood. Back-cross them to the strain used for injection and sort positive F1 embryos (according to the expression of the transgenesis marker). F1 embryos can be analyzed phenotypically or raised to adulthood.
***Note:*** *depending on the transgenesis marker used, it may be possible to evaluate the level of mosaicism in injected embryos. In that case, injected embryos could be further sorted according to their level of mosaicism.*

### References


Ablain, J., Durand, E.M., Yang, S., Zhou, Y., Zon, L.I., 2015. A CRISPR/Cas9 vector system for tissue-specific gene disruption in zebrafish. Developmental cell 32, 756-764.

Campbell, P.D., Heim, A.E., Smith, M.Z., Marlow, F.L., 2015. Kinesin-1 interacts with Bucky ball to form germ cells and is required to pattern the zebrafish body axis. Development 142, 2996-3008.

Gagnon, J.A., Valen, E., Thyme, S.B., Huang, P., Ahkmetova, L., Pauli, A., Montague, T.G., Zimmerman, S., Richter, C., Schier, A.F., 2014a. Efficient mutagenesis by Cas9 protein-mediated oligonucleotide insertion and large-scale assessment of single-guide RNAs. PloS one 9, e98186.

Gagnon, J.A., Valen, E., Thyme, S.B., Huang, P., Akhmetova, L., Pauli, A., Montague, T.G., Zimmerman, S., Richter, C., Schier, A.F., 2014b. Efficient mutagenesis by Cas9 protein-mediated oligonucleotide insertion and large-scale assessment of single-guide RNAs. PloS one 9, e98186.

Hartley, J.L., Temple, G.F., Brasch, M.A., 2000. DNA cloning using in vitro site-specific recombination. Genome research 10, 1788-1795.

Heim, A.E., Hartung, O., Rothhamel, S., Ferreira, E., Jenny, A., Marlow, F.L., 2014. Oocyte polarity requires a Bucky ball-dependent feedback amplification loop. Development (Cambridge, England) 141, 842-854.

Hsu, P.D., Scott, D.A., Weinstein, J.A., Ran, F.A., Konermann, S., Agarwala, V., Li, Y., Fine, E.J., Wu, X., Shalem, O., Cradick, T.J., Marraffini, L.A., Bao, G., Zhang, F., 2013. DNA targeting specificity of RNA-guided Cas9 nucleases. Nat Biotechnol 31, 827-832.

Hwang, W.Y., Fu, Y., Reyon, D., Maeder, M.L., Tsai, S.Q., Sander, J.D., Peterson, R.T., Yeh, J.R., Joung, J.K., 2013. Efficient genome editing in zebrafish using a CRISPR-Cas system. Nature biotechnology 31, 227-229.

Kaufman, O.H., Lee, K., Martin, M., Rothhamel, S., Marlow, F.L., 2018. rbpms2 functions in Balbiani body architecture and ovary fate. PLoS genetics 14, e1007489.

Kawakami, K., Takeda, H., Kawakami, N., Kobayashi, M., Matsuda, N., Mishina, M., 2004. A transposon-mediated gene trap approach identifies developmentally regulated genes in zebrafish. Developmental cell 7, 133-144.

Kim, H.J., Lee, H.J., Kim, H., Cho, S.W., Kim, J.S., 2009. Targeted genome editing in human cells with zinc finger nucleases constructed via modular assembly. Genome Res 19, 1279-1288.

Kwan, K.M., Fujimoto, E., Grabher, C., Mangum, B.D., Hardy, M.E., Campbell, D.S., Parant, J.M., Yost, H.J., Kanki, J.P., Chien, C.B., 2007. The Tol2kit: a multisite gateway-based construction kit for Tol2 transposon transgenesis constructs. Developmental dynamics : an official publication of the American Association of Anatomists 236, 3088-3099.

Leu, D.H., Draper, B.W., 2010. The ziwi promoter drives germline-specific gene expression in zebrafish. Developmental dynamics : an official publication of the American Association of Anatomists 239, 2714-2721.

Meeker, N.D., Hutchinson, S.A., Ho, L., Trede, N.S., 2007. Method for isolation of PCR-ready genomic DNA from zebrafish tissues. Biotechniques 43, 610, 612, 614.

Montague, T.G., Cruz, J.M., Gagnon, J.A., Church, G.M., Valen, E., 2014. CHOPCHOP: a CRISPR/Cas9 and TALEN web tool for genome editing. Nucleic acids research 42, W401-407.
